# Regional astrocyte interferon-γ signaling regulates immunoproteasome-mediated protection during chronic autoimmunity

**DOI:** 10.1101/2019.12.23.887455

**Authors:** Brandon C. Smith, Maksim Sinyuk, Julius E. Jenkins, Morgan W. Psenicka, Jessica L. Williams

## Abstract

In early autoimmune neuroinflammation, interferon (IFN)γ and its upregulation of the immunoproteasome (iP) is pathologic. However, during chronic multiple sclerosis (MS), IFNγ has protective properties and, the role of the iP remains to be fully elucidated. Here, we demonstrated that IFNγ signaling in regional astrocytes induces the iP and protects the CNS during autoimmunity. In MS tissue, iP expression was enhanced in spinal cord compared to brainstem lesions, which correlated with a decrease in oxidative stress. *In vitro*, IFNγ stimulation enhanced iP expression, reduced reactive oxygen species burden, and decreased oxidatively damaged and poly-ubiquitinated protein accumulation preferentially in human spinal cord astrocytes, which was abrogated with the use of the iP inhibitor, ONX 0914. During the chronic phase of an MS animal model, experimental autoimmune encephalomyelitis (EAE), ONX 0914 treatment exacerbated disease and led to increased oxidative stress and poly-ubiquitinated protein build-up. Finally, mice with astrocyte-specific loss of the IFNγ receptor exhibited worsened chronic EAE associated with reduced iP expression, enhanced lesion size and oxidative stress and poly-ubiquitinated protein accumulation in astrocytes. Taken together, our data reveal a protective role for IFNγ in chronic neuroinflammation and identify a novel function of the iP in astrocytes during CNS autoimmunity.

## Introduction

Multiple sclerosis (MS) is the most common chronic inflammatory and neurodegenerative disease of the central nervous system (CNS) (1). During the pathogenesis of MS, there is immune cell infiltration, demyelination, and reactive gliosis within CNS lesions in multiple regions (2). Relapsing-remitting MS (RRMS) is a subtype that affects approximately 85% of patients and is characterized by episodic periods of neurological dysfunction, often associated with inflammation, followed by partial or complete recovery. A significant proportion of these patients go on to develop secondary progressive MS (SPMS), during which they have fewer remissions and increasing atrophy, correlating with progressive disability (3, 4). Primary progressive (PPMS) a third subtype of MS, affects approximately 15% of patients and is associated with continuous, progressive loss of neurological function after initial diagnosis, without periods of remission (5, 6). Of note, it is thought that the pathology associated with RRMS has a relatively significant inflammatory component, while in SPMS and PPMS, inflammation is relatively limited (6). Thus, it is not surprising that the 12 immunomodulatory, FDA-approved therapies for RRMS (7) have limited effectiveness in SPMS and PPMS patients (8-10) and in some cases have resulted in patient worsening (11, 12). Indeed, neutralizing specific inflammatory cytokines in MS patients resulted in exacerbated neurological deficits (13) without reducing lesion load (14), suggesting a complex, and potentially protective role for inflammatory cytokines during chronic MS that is not fully understood.

Astrocytes are the most numerous cell type in the CNS and are found in and around MS lesions. Although they have long been considered bystanders of MS pathology, their role in the initiation and resolution of disease is becoming more appreciated (15, 16). During MS and experimental autoimmune encephalomyelitis (EAE), an animal model of MS, astrocytes are known to exhibit regional heterogeneity in gene expression and response to inflammation (17-19). Indeed, ablation of astrocytes following several types of CNS injury leads to sustained inflammation, impaired repair, and increased neurodegeneration (20-29), suggesting that a diverse astrocytic response is critical in healthy tissue preservation and support, minimizing CNS bystander damage during neuroinflammation. However, during chronic MS, inflammation is largely overlooked as it is thought to have a lesser role, especially in SPMS and PPMS patients. Thus, relatively little is known about how regionally distinct astrocytes respond to chronic inflammation to facilitate damage control and recovery.

The role of the cytokine interferon (IFN)γ in MS and EAE has been a paradox for more than 3 decades. Many early studies describe a solely proinflammatory and pathologic function in disease (30, 31). However, more recent evidence supports additional protective roles, particularly in chronic stages, suggesting IFNγ has complex, stage-dependent pleiotropic effects in MS and EAE (32-39). In progressive MS patients, improving symptoms correlated with high levels of serum IFNγ, while patients with clinical worsening had relatively low levels of serum IFNγ (37). In EAE, systemic or intraventricular administration of IFNγ in mice and marmosets during chronic phases reduced disease severity, demyelination and mortality (40-42), and significantly delayed relapses in a murine model of chronic-relapsing EAE (43). Likewise, neutralizing IFNγ exacerbated disease and made an EAE-resistant mouse strain susceptible (40, 41, 43-46). These studies were corroborated by eliminating IFNγ signaling using genetic deletion of *Ifng* or *Ifngr1*, which resulted in higher susceptibly to EAE, increased incidence, more extensive inflammation, encompassing both the spinal cord and hindbrain, and exacerbated disease compared to WT animals (47-55). Further, only mice with intact IFNγ signaling were able to recover from EAE (56). Using a signaling deficient dominant-negative *Ifngr1* driven by the *Gfap* promotor, the protective effects of IFNγ signaling during chronic EAE was linked to astrocytes (57). While follow-up studies indicate these effects may be due to altered cytokine release influencing microglia (58), the astrocyte-specific mechanisms of IFNγ-mediated protection are not defined.

Our study demonstrating that IFNγ signaling in regional astrocytes mediates protection during chronic autoimmune neuroinflammation through induction of the immunoproteasome (iP) has clear clinical applications. Inefficacy of first line immunomodulatory therapies in chronic-progressive MS patients demands the identification of novel therapeutic approaches. Since the iP has a critical role in T cell activation during the early, inflammatory phases of MS and EAE, which is thought to drive pathogenesis, and proteasome inhibitors have recently been utilized for the treatment of certain types of cancer, proteasome inhibitors are being discussed as a potential treatment strategy for MS patients (59, 60). Given that the role of the iP in the CNS is largely unexplored and may have a neuroprotective function in astrocytes, further understanding of the iP is of great importance and incredibly timely (61). Our study suggests that the iP is a potential mediator of protection during chronic CNS autoimmunity following astrocyte IFNγ signaling and identification of an endogenous inhibitor of the iP might represent a novel therapeutic target that would benefit chronic MS patients with specific patterns of neuroinflammation. In summary, these findings advance our understanding of the astrocyte adaptive immune response during chronic CNS autoimmunity, identify regionally distinct protective role for astrocytes, and suggest that defining upstream targets that modulate iP expression would facilitate the identification of new targets for the treatment of SPMS and PPMS patients.

## Results

In several neurodegenerative diseases, the iP is known to clear reactive oxygen species (ROS) and degrade poly-ubiquitinated proteins (62-64); and, there is evidence that the iP may be dysregulated during MS and experimental autoimmune encephalomyelitis (EAE), an animal model of MS (64-66). Additionally, astrocytes have been implicated in modulating neuroinflammation and neurodegeneration (67-69); however, the role of the astrocyte iP during MS and EAE has not yet been fully elucidated. To determine the role of the iP in MS brain, we evaluated the mRNA expression of genes associated with the classical proteasome (*PSMB7*) and the iP (*PSMB8*) in previously published MS patient tissue microarray data (70). Strikingly, we found *PSMB7* to be significantly downregulated while *PSMB8* was significantly increased in white matter lesions (WML) compared to controls and normal appearing white matter (NAWM) (Figure 1, A and B). Although lesions in the brainstem and spinal cord are common and result in pronounced disability during MS (71, 72) and EAE (73), relatively little is known about lesion pathology in these regions. Thus, we examined NAWM and areas of demyelination in the brainstem (Figure 1C) and spinal cord (Figure 1D) of MS patients, labeling an iP subunit (LMP7) and an indicator of oxidative stress (PRDX6) within astrocytes by immunohistochemistry (IHC). In WMLs, there was an inverse relationship between LMP7 and PRDX6 specifically in the spinal cord (Figure 1E), suggesting that the iP may play a role in reducing oxidative stress in MS lesions and that there may be a difference in iP expression and/or function in regional astrocytes.

**Figure 1.**
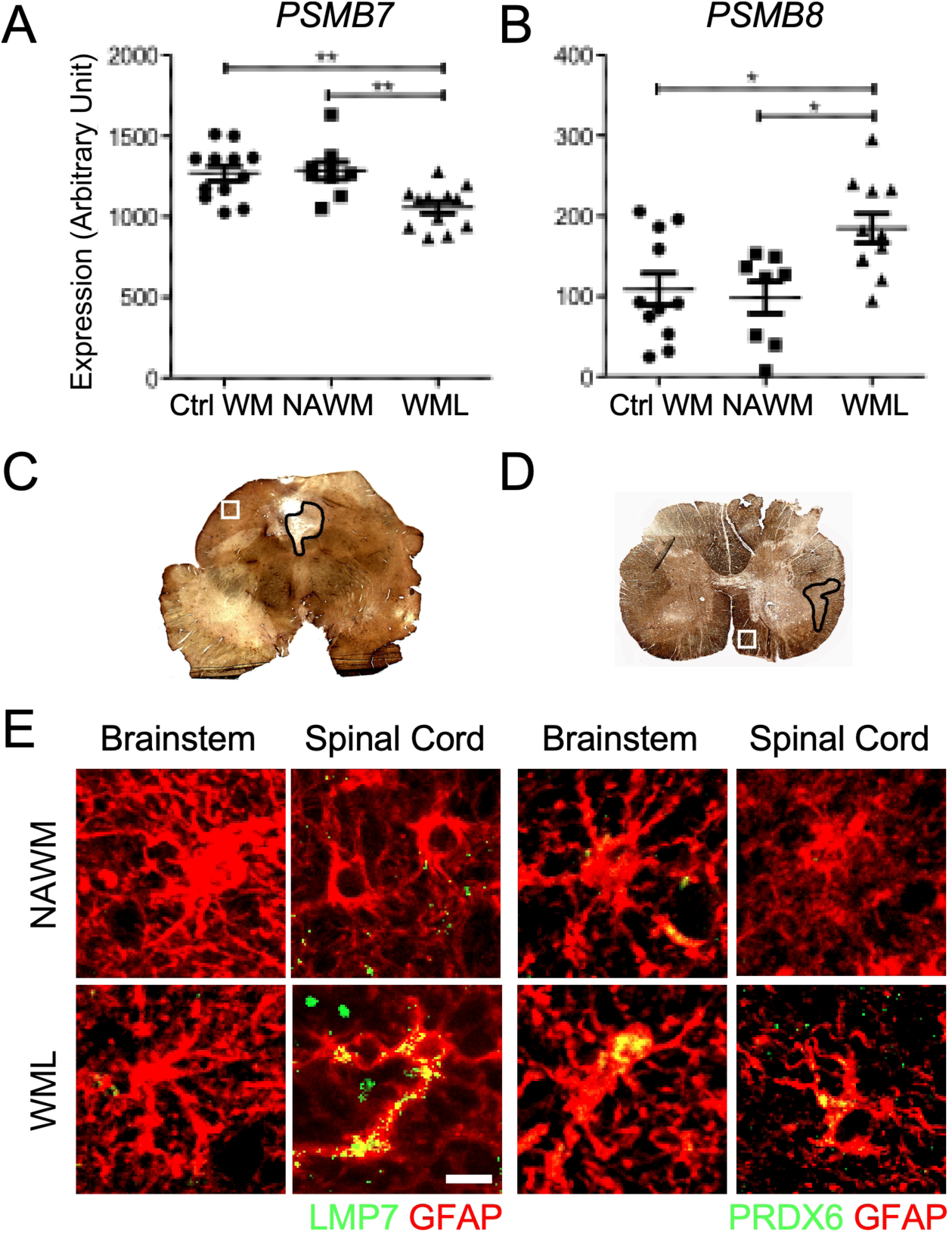
Regional astrocyte oxidative stress and iP expression in chronic MS WMLs. (**A** and **B**) RNA was extracted from the cortical WM of control and MS (*n*=7-12) patients and analyzed via microarray and the relative expression of (**A**) *PSMB7* and (**B**) *PSMB8* were assessed as described (70). (**C** and **D**) MS demyelinating lesions (black outline) and NAWM (white square) were identified by PLP labeling of the (**C**) brainstem and (**D**) spinal cord in MS patients. (**E**) GFAP^+^ astrocytes (red) were co-labeled with either LMP7 or PRDX6 (green) in NAWM or in WML of the brainstem or spinal cord. Scale bar: 10 μM. \**P* < 0.05, \*\**P* < 0.01. Ctrl WM, control white matter; NAWM, normal appearing white matter; WML, white matter lesion.

Astrocytes are regionally heterogeneous in morphology, gene expression, and function during physiological and pathological conditions (74, 75). Since the iP is readily induced by IFNγ, which is present at all stages of MS (34, 76), and astrocytes are in and around MS lesions and respond to IFNγ, we further explored heterogenous regulation of the astrocyte iP between the brainstem and spinal cord driven by IFNγ. Using primary human astrocytes from the brainstem and spinal cord (Supplemental Figure 1), we analyzed iP transcript levels following IFNγ stimulation over the course of 48 h. Expression levels of the iP subunits *PSMB8, PSMB9*, and *PSMB10* increased in spinal cord astrocytes compared to those in brainstem-derived astrocytes, with the most robust transcript upregulation occurring at 48 h (Figure 2A). Of note, there was no appreciable change in constitutive proteasome subunit transcript in either region following IFNγ stimulation (Supplemental Figure 2). We confirmed regional differences in iP expression in astrocytes at the protein level by analyzing protein lysate from human brainstem and spinal cord astrocytes incubated with and without IFNγ (Figure 2, B and C). Although IFNγ expression is robust and perpetuates acute inflammation in both MS and EAE, it is still present during chronic disease, albeit at lower levels (77, 78). To determine the regional sensitivity of astrocytes to IFNγ stimulation, we measured the transcript expression levels of iP subunits following an IFNγ dose titration over 24 or 48 h. Interestingly, compared to those from the brainstem, all of the iP subunits were upregulated specifically in astrocytes from the spinal cord following exposure to low concentrations of IFNγ after 48 h (Figure 2D). To determine if regional differences in astrocyte iP expression were present *in vivo* during neuroinflammation, we induced EAE in *Ifngr1*^fl/fl^ *Tie2*-Cre^+^ mice, in which the IFNγ receptor (IFNGR1) is deleted from endothelial cells of the blood-brain barrier (79, 80), conferring inflammation in both the brainstem and spinal cord and maintaining genetically WT astrocytes. Signs of both classical EAE, primarily affecting the spinal cord, and atypical EAE, which affects the hindbrain, was monitored (Supplemental Figure 3A) (81, 82). To determine if iP expression is regionally distinct, we first confirmed that lesion size between the brainstem and spinal cord was consistent in *Ifngr1*^fl/fl^ *Tie2*-Cre^+^ mice (Supplemental Figure 3B). While the average lesion size was similar between regions in *Ifngr1*^fl/fl^ *Tie2*-Cre^+^ mice, iP expression in spinal cord astrocytes was significantly increased compared to those in the brainstem (Supplemental Figure 3C). These data demonstrate that both in human astrocytes and in an *in vivo* model of regional autoimmune neuroinflammation, astrocyte iP expression is preferentially upregulated by IFNγ in spinal cord astrocytes compared to those in the brainstem. This suggests that in the spinal cord, the iP is potentially a primary astrocyte-mediated protection mechanism during neuroinflammation and that brainstem astrocytes may engage alternate pathways.

**Figure 2.**
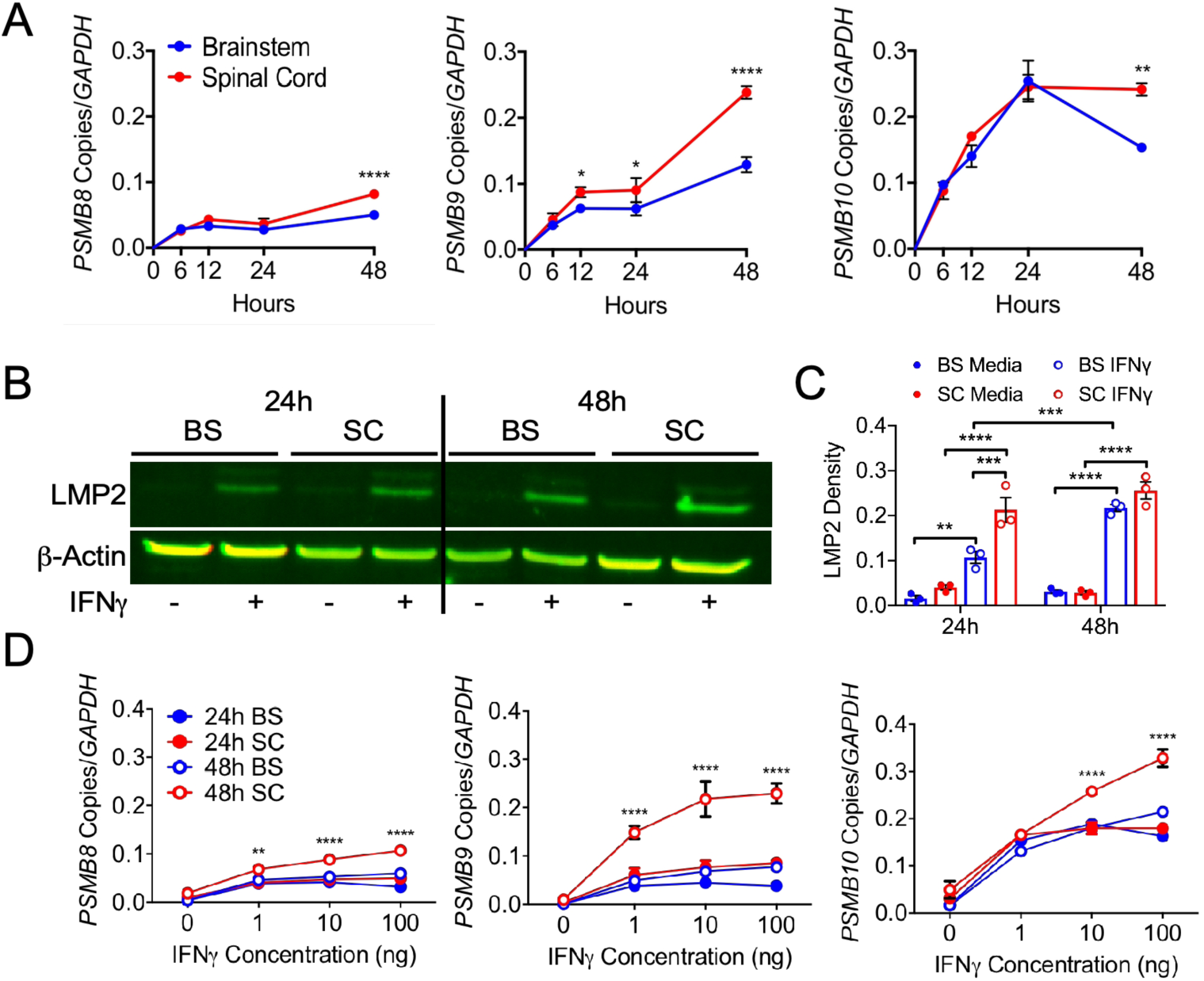
IFNγ-regulated expression of the iP in regional human astrocytes. (**A**) Human brainstem and spinal cord astrocytes were stimulated with 10 ng/ml IFNγ for 0, 6, 12, 24, or 48 h and RNA was collected and analyzed for transcript levels of *PSMB8, PSMB9*, and *PSMB10* by qRT-PCR, normalized to copies of *GAPDH*. (**B**) Human brainstem and spinal cord astrocytes were stimulated with or without 10 ng/ml IFNγ for 24 or 48 h and protein lysate was assessed for levels of LMP2, normalizing to β-actin. (**C**) Human brainstem and spinal cord astrocytes were stimulated with 0, 1, 10, or 100 ng/ml IFNγ for 24 or 48 h and RNA was collected and analyzed for transcript levels of *PSMB8, PSMB9*, and *PSMB10* by qRT-PCR, normalized to copies of *GAPDH*. Data represent the mean ± SEM from 3 independent experiments. \**P* < 0.05, \*\**P* < 0.01, \*\*\**P* < 0.001, \*\*\*\**P* < 0.0001 by 2-way ANOVA. BS, brainstem; SC, spinal cord.

**Figure 3.**
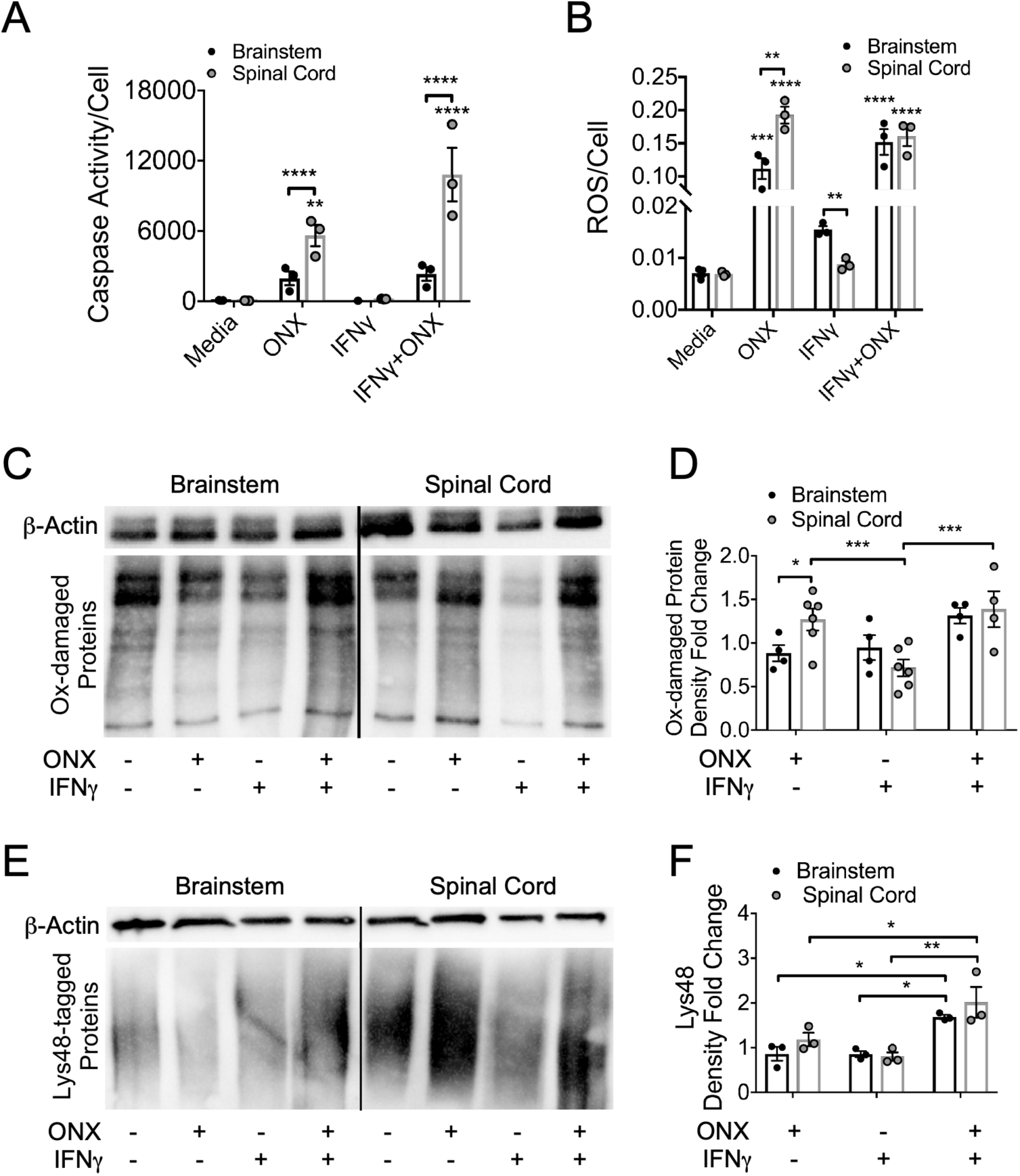
The effect of iP inhibition on regional astrocyte death, ROS production, oxidatively damaged and poly-ubiquitinated protein load. (**A-E**) Human brainstem and spinal cord astrocytes were stimulated with or without 10 ng/ml IFNγ and/or ONX 0914 for 48 h. (**A**) Caspase activity and (**B**) ROS were measured and normalized to live cells. (**C**) Protein lysate was collected, derivatized, and (**D**) oxidatively damaged protein accumulation was quantified and normalized to β-actin levels. (**E**) Protein lysate was probed for Lys48-labeled poly-ubiquitinated proteins, normalized to β-actin expression, (**F**) and quantified. Data represent the mean ± SEM from 3 independent experiments. \**P* < 0.05, \*\**P* < 0.01, \*\*\**P* < 0.001, \*\*\*\**P* < 0.0001 by 2-way ANOVA. ROS, reactive oxygen species; Ox-damaged, oxidatively damaged.

The best known function of the iP is its role in antigen processing; however, since astrocytes present little or no antigen (83-85), alternate functions of the iP were examined, namely, clearance of ROS and oxidatively damaged and poly-ubiquitinated proteins (86). To determine the role of the iP in astrocyte viability and ROS clearance, we treated regional human astrocytes with or without IFNγ and a specific inhibitor of the iP, ONX 0914 (87-89). Following iP inhibition, an increase in caspase activity was observed only in spinal cord astrocytes compared to media treatment, which was significantly enhanced compared to those from the brainstem (Figure 3A). Next, we observed an overall increase in cellular ROS in astrocytes treated with ONX 0914, with enhanced ROS in spinal cord astrocytes compared to brainstem astrocytes. Interestingly, in cells treated only with IFNγ, astrocytes from the spinal cord produced less ROS than brainstem-derived astrocytes (Figure 3B). Assessment of oxidative protein damage revealed an increase in spinal cord versus brainstem astrocytes following iP inhibition. There was also a treatment effect in spinal cord astrocytes with a reduction in oxidatively damaged proteins following IFNγ treatment (Figure 3, C and D). Assessment of poly-ubiquitinated proteins revealed an increase in both spinal cord and brainstem astrocytes following iP inhibition in the presence of IFNγ (Figure 3, E and F). Taken together, these data suggest that IFNγ-mediated induction of the iP in spinal cord astrocytes reduces ROS and oxidatively damaged and poly-ubiquitinated proteins which may lead to enhanced cell survival.

Although iP inhibition during acute EAE resulted in ameliorated disease associated primarily with dampened peripheral immune responses (90), the role of the iP during the relatively less inflammatory chronic phase of EAE has not yet been explored. We induced EAE in WT C57Bl/6 mice, which develop spinal cord-associated inflammation, and two days after peak disease, we administered a single dose of ONX 0914 or vehicle and the clinical course was monitored. Following treatment, EAE rapidly worsened in mice receiving ONX 0914 compared to vehicle-treated mice (Figure 4A). Further, IHC labeling revealed an increase in lesion size and a decrease in myelin in ONX 0914-treated mice (Figure 4, B and C). To determine if this corresponded to a reduction in oxidative stress and accumulation of poly-ubiquitinated protein accumulation, we performed IHC analysis on tissue 7 days following treatment. Indeed, IHC labeling of PRDX6 indicated enhanced oxidative stress and oxidative stress within GFAP^+^ astrocytes in lesions of ONX 0914-treated compared to vehicle-treated mice (Figure 4, D and E). Similarly, Lys48 labeling revealed an increase in total poly-ubiquitination and poly-ubiquitination within GFAP^+^ astrocytes in lesions of ONX 0914-treated mice compared to those treated with vehicle (Figure 4, F and G). These data suggest that the iP has a role in reducing oxidative stress and poly-ubiquitinated proteins in astrocytes during chronic EAE, which may contribute to reduced lesion size and disease recovery.

**Figure 4.**
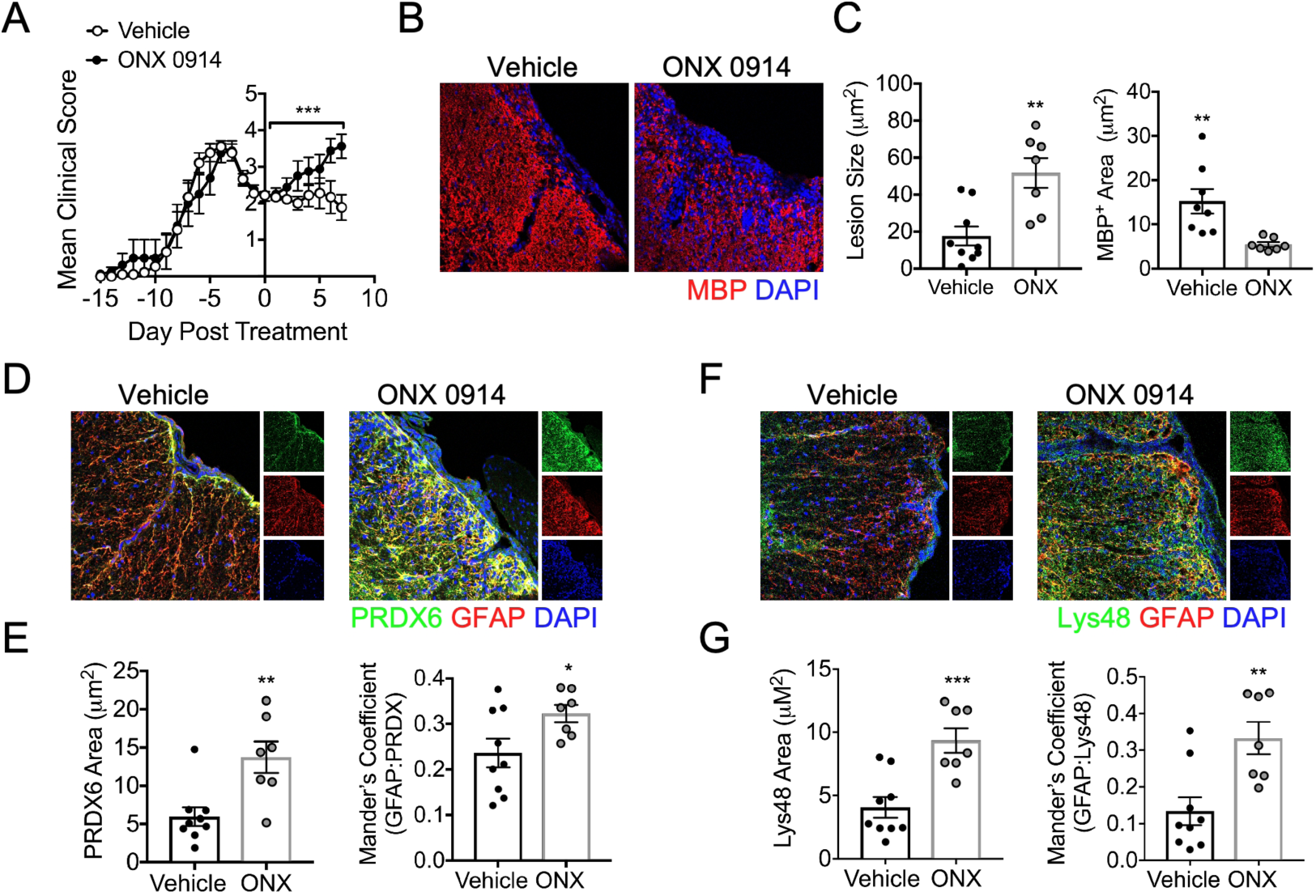
iP inhibition exacerbates chronic EAE. Animals were randomly assigned to a treatment group and EAE was induced in WT C57Bl/6 mice. After a stabilization or reduction in EAE clinical score was observed for two consecutive days, 10 mg/kg ONX 0914 (*n* = 9) or vehicle (*n* = 7) was administered to mice i.p. (**A**) Following treatment, EAE clinical course was monitored in a blinded fashion and data is presented relative to day post treatment. Seven days after vehicle or ONX 0914 treatment, mice were perfused and CNS tissue was prepared for IHC analysis and ventral white matter tracts of the lumbar spinal cord were imaged using confocal microscopy at 20x magnification. (**B**) Tissue sections were labeled for MBP (red) and nuclei were counterstained with DAPI (blue). (**C**) Lesion area and MBP^+^ area were quantified using ImageJ software. **(D)** Tissue sections were labeled for PRDX6 (green), GFAP (red), and nuclei were counterstained with DAPI (blue). **(E)** Total PRDX6 area and PRDX6 colocalized with GFAP were analyzed. **(F)** Tissue sections were labeled for Lys48 (green), GFAP (red), and nuclei were counterstained with DAPI (blue). **(G)** Total Lys48 area and Lys48 colocalized with GFAP were analyzed. Data in **A** represent the mean ± SEM combined from 2 independent experiments and were analyzed by Mann-Whitney *U* test for nonparametric data. Data in **B-G** represent the mean ± SEM combined from 2 independent experiments and were analyzed by 2-tailed Student’s *t* test. \**P* < 0.05, \*\**P* < 0.01, \*\*\**P* < 0.001.

Diminished IFNγ signaling in astrocytes is known to exacerbate chronic EAE (81). We demonstrated that the iP is regulated by IFNγ-signaling and we observed an increase in oxidative stress and poly-ubiquitinated protein accumulation specifically in astrocytes during *in vivo* iP inhibition. To further examine the role of IFNγ-mediated iP induction in astrocytes during EAE, we immunized mice in which *Gfap*-expressing astrocytes are deficient in *Ifngr1* (Supplemental Figure 4). Following EAE onset, *Ifngr1*^fl/fl^ *Gfap-*Cre^+^ and *Ifngr1*^fl/fl^ littermate control mice exhibited similar acute disease; however, chronic disease was significantly exacerbated in *Ifngr1*^fl/fl^ *Gfap-*Cre^+^ mice and weight loss was enhanced (Figure 5, A and B). Further, compared to *Ifngr1*^fl/fl^ controls, *Ifngr1*^fl/fl^ *Gfap-*Cre^+^ mice had increased lesion size and reduced myelin basic protein (MBP) expression by IHC (Figure 5, C and D). To determine if there was a corresponding reduction in astrocyte iP expression and function in *Ifngr1*^fl/fl^ *Gfap-*Cre^+^ mice, we performed IHC analysis during chronic EAE. As expected, there was a decrease in both total iP expression and iP colocalization with GFAP^+^ astrocytes within the lesions of *Ifngr1*^fl/fl^ *Gfap-*Cre^+^ compared to *Ifngr1*^fl/fl^ mice (Figure 5, E and F). Further IHC labeling revealed an increase in both total PRDX6 and Lys48 and PRDX6 and Lys48 colocalized with GFAP^+^ astrocytes within the lesions of *Ifngr1*^fl/fl^ *Gfap-*Cre^+^ compared to *Ifngr1*^fl/fl^ mice (Figure 5, G and H), suggesting enhanced oxidative stress and poly-ubiquitinated protein accumulation specifically in astrocytes of *Ifngr1*^fl/fl^ *Gfap-*Cre^+^ mice (Figure 5, I and J). These data suggest that during chronic EAE, IFNγ signaling in astrocytes enhances iP expression, which leads to a reduction in CNS tissue damage, allowing recovery from clinical disease.

**Figure 5.**
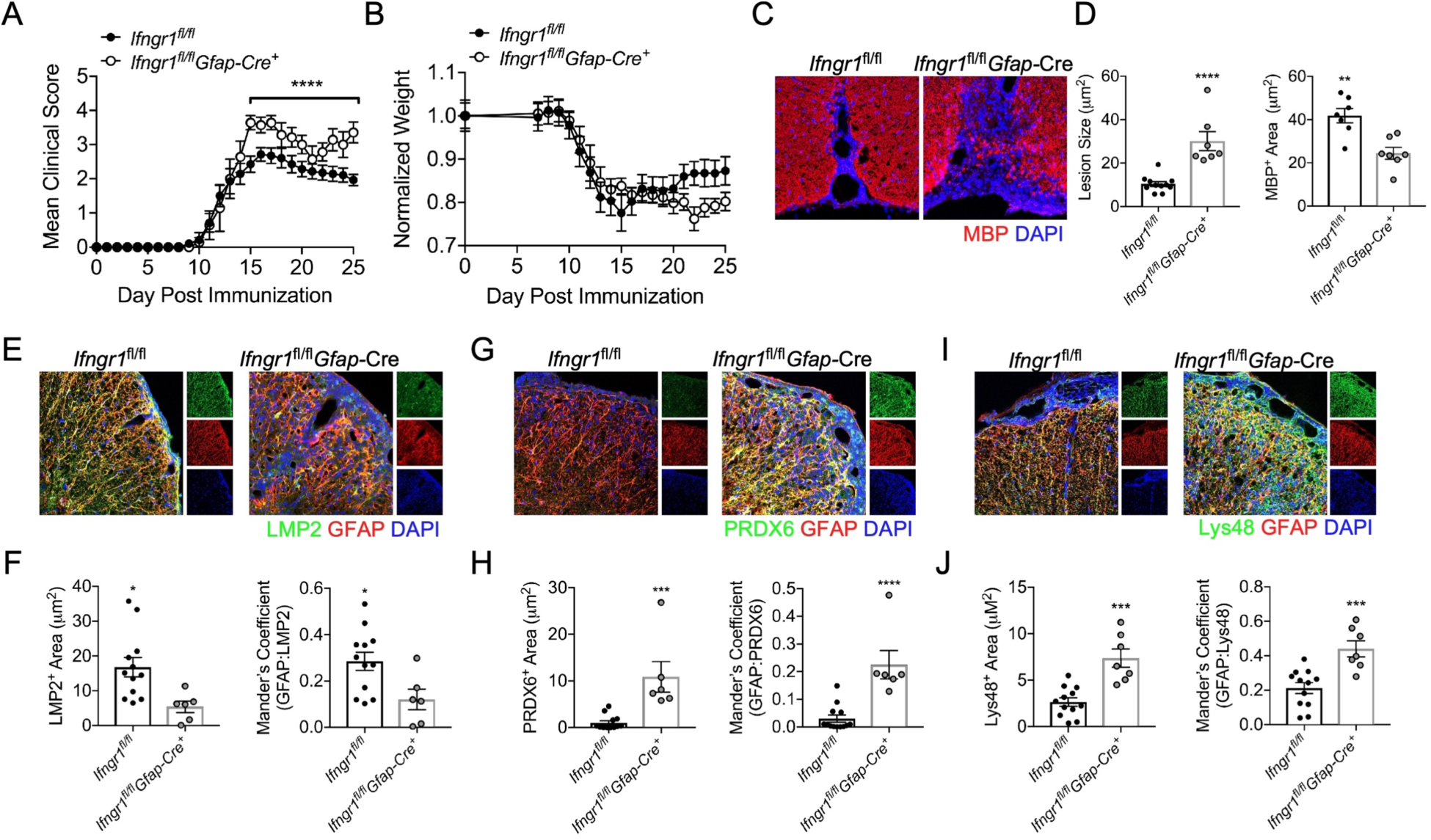
IFNγ signaling upregulates the iP in astrocytes during EAE. EAE was induced in *Ifngr1*^fl/fl^ *Gfap*-Cre^+^ mice (*n* = 7) and *Ifngr1*^fl/fl^ littermates (*n* = 12) and **(A)** EAE clinical course and **(B)** weight loss were blindly monitored. Following 25 days post-immunization, mice were perfused and the CNS was removed and cryopreserved for IHC analysis. Ventral white matter tracts of the lumbar spinal cord were imaged using confocal microscopy at 20x magnification. (**C**) Tissue sections were labeled for MBP (red) and nuclei were counterstained with DAPI (blue). (**D**) Lesion area and MBP^+^ area were quantified using ImageJ software. **(E)** Tissue sections were labeled for LMP2 (green), GFAP (red), and nuclei were counterstained with DAPI (blue). **(F)** Total LMP2 area and LMP2 colocalized with GFAP were analyzed. **(G)** Tissue sections were labeled for PRDX6 (green), GFAP (red), and nuclei were counterstained with DAPI (blue). **(H)** Total PRDX6 area and PRDX6 colocalized with GFAP were analyzed. **(I)** Tissue sections were labeled for Lys48 (green), GFAP (red), and nuclei were counterstained with DAPI (blue). **(J)** Total Lys48 area and Lys48 colocalized with GFAP were analyzed. Data in **A** represent the mean ± SEM combined from 2 independent experiments and were analyzed by Mann-Whitney *U* test for nonparametric data. Data in **C-J** represent the mean ± SEM combined from 2 independent experiments and were analyzed by 2-tailed Student’s *t* test. \**P* < 0.05, \*\**P* < 0.01, \*\*\**P* < 0.001, \*\*\*\**P* < 0.0001.

## Discussion

Interferonγ is primarily pro-inflammatory in acute stages of MS and EAE (78); however, there is evidence to suggest that in chronic MS and EAE, IFNγ has protective functions (37, 38, 81). In this study, we confirm these findings and demonstrate that IFNγ signaling in astrocytes leads to the upregulation of the iP, which dampens chronic EAE severity. Our studies indicate that IFNγ-mediated upregulation of the iP leads to preservation of astrocyte integrity by reducing ROS, degrading oxidatively damaged and poly-ubiquitinated proteins. *In vivo*, IFNγR signaling was critical for iP upregulation in astrocytes, which was associated with a reduction in lesion size and disease severity during chronic EAE. Further, studies using postmortem MS tissue, *in vitro* astrocytes, and models of EAE showed enhanced iP expression in the spinal cord compared to the brainstem, suggesting that spinal cord astrocytes more readily upregulate the iP upon IFNγ stimulation, leading to enhanced region-specific protection during CNS autoimmunity.

These findings reveal the importance of IFNγ signaling and its role in astrocyte iP regulation during chronic stages of neurodegenerative disease, and they highlight the stage-specific roles of IFNγ during MS and EAE. Indeed, previous studies have shown that iP inhibition during acute disease leads to EAE amelioration by dampening peripheral immune responses (90, 91). Importantly, it has recently been shown that IFNγ diverts OPCs from differentiating into mature oligodendrocytes to cells with antigen presenting capabilities via upregulation of the iP, a critical immune-priming process during acute EAE (92). However, during chronic EAE, IFNγR signaling specifically in astrocytes was beneficial, resulting in smaller lesions, less demyelination and a reduction in EAE severity in control mice compared to those with astrocytes deficient in IFNγR (81). Here, we extend those findings by demonstrating that upregulation of the iP is one potential mechanism by which IFNγ signaling protects the CNS during chronic neuroinflammation. We demonstrate that the iP is expressed in astrocytes during chronic MS and that administration of a specific iP inhibitor during chronic EAE exacerbates clinical disease, likely as a result of inhibiting the alternate functions of the iP including clearance of ROS and poly-ubiquitinated proteins.

While IFNγ signaling is known to directly induce iP expression (93), other factors including aging, stress, and ROS also contribute to iP upregulation (94-96). Due to normal metabolic processes and oxidative and age-related stress, there is a basal level of iP expression that supports cellular homeostasis (94). Without a functioning iP, damaged proteins can rapidly accumulate under oxidative stress conditions, like EAE and MS, to an extent that exceeds the proteolytic capacity of the constitutive proteasome, which leads to the formation of harmful protein aggregates and cell apoptosis (97-99). Importantly, oxidative modifications are elevated during EAE (99, 100) and accumulate primarily in astrocytes in MS lesions (101, 102). Consistent with this, oxidized and polyubiquitinated proteins accumulated in chronic MS white matter with reduced iP peptidase activity (103) and in the brains of mice lacking the iP subunits LMP2 and LMP7 (97, 104). Further, LMP7-deficient mice had more severe CNS oxidative damage and exacerbated EAE (97). While the pathological consequences of impaired iP activity in chronic MS and EAE are unknown, the impact of accumulated, damaged protein aggregates and reduced degradation of various signaling and pro-apoptotic molecules likely contribute to neurodegeneration.

These findings broaden our understanding of astrocyte heterogeneity in the CNS. Although regional heterogeneity of astrocytes in neurophysiologic functions (18, 19, 105) and in many disease models (106-108), including EAE (75, 109, 110), has been appreciated, there are no studies to date that have described a regional role for the iP in the CNS. However, a regional difference in Type I IFN signaling in astrocytes has been described (111). Here, we expand on those findings, demonstrating a regional increase in iP expression preferentially in spinal cord astrocytes compared to those from the brainstem in response to IFNγ. We show that this increase in iP expression in spinal cord astrocytes is protective, as inhibition of the iP results in reduced astrocyte viability, increased ROS production, and accumulation of damaged proteins. This diverse astrocytic response is critical in healthy tissue preservation and support as it minimizes prolonged CNS exposure to cytotoxic inflammation. Indeed, ablation of astrocytes following several types of CNS injury leads to sustained inflammation, impaired blood-brain barrier repair, and increased neurodegeneration (20-29). Thus, the survival of astrocytes following damage is key in CNS recovery. Taken together, our data suggests that the astrocyte iP may be a key mechanism of protection in the spinal cord while other distinct CNS regions may rely on alternate pathways to facilitate damage control and recovery during chronic neurodegeneration and CNS inflammation.

Our study demonstrating a role for the astrocyte iP in promoting protection during chronic neuroinflammation has definite clinical implications. It has been proposed that inhibition of the iP can serve as an effective therapeutic modality for MS; however, given the data presented here, consideration of disease stage is of utmost importance (112, 113). In progressive MS patients, improving symptoms correlated with high levels of serum IFNγ, while patients with clinical worsening had relatively low levels of serum IFNγ (37), suggesting IFNγ-induced iP expression may have a role in MS stabilization or even recovery. Since the iP is critical for maintenance of astrocytes and recovery from chronic autoimmunity, inhibition would likely greatly exacerbate patient symptoms if given during chronic stages of MS.

## Methods

### Human tissue IHC

Brainstem and spinal cord tissue from MS patients were collected according to the established rapid autopsy protocol approved by the Cleveland Clinic Institutional Review Board (114). Tissue was removed, fixed in 4% paraformaldehyde and sectioned. Demyelinated lesions were identified by immunostaining with proteolipid protein (PLP) as described previously (115, 116) and followed by collection of subsequent sections for immunoproteasome and astrocyte labeling. Antigen retrieval was performed using 10 μM citrate buffer and boiling briefly. Sections were blocked with 3% goat serum and 0.01% Triton X-100 (Sigma-Aldrich) for 1 h at room temperature and then exposed to antibodies specific for human LMP7 (ThermoFisher Scientific; MA5-15890) or PRDX6 (Abcam; ab59543) and GFAP (Invitrogen; 13-0300) for 4 days at 4°C. Sections were then washed with PBS-Tween 20 and secondary antibodies conjugated to Alexa Fluor 488 or 555 (ThermoFisher Scientific) were applied for 1 h at room temperature. Sections were then treated with 0.3% sudan black in 70% ethanol for 3 min, analyzed using the 20x objective of a confocal microscope LSM 800 (Carl Zeiss), and analyzed using ImageJ (NIH).

### Human astrocytes

Primary adult human brain stem and spinal cord astrocytes were obtained from ScienCell and grown according to provided protocols in complete ScienCell Astrocyte Medium. Briefly, primary human astrocytes were isolated from normal brain stem or spinal cord tissue and at P0 were tested for morphology by phase contrast and relief contrast microscopy and GFAP positivity by immunofluorescence. Cell number, viability (≥ 70%), and proliferative potential (≥ 15 pd) were also assessed and negative screening for potential biological contaminants was confirmed prior to cryopreservation and receipt of frozen cells at P1. Purity was determined by qRT-PCR (Supplemental Figure 1).

### EAE induction

Animals of mixed sex were induced for EAE at 8-10 weeks of age. *Gfap*-Cre^+^ line 77.6, *Tie2*-Cre^+^, and *Ifngr1*^fl/fl^ mice were obtained commercially from The Jackson Laboratory and housed under specific pathogen-free conditions. Mice were crossed according to standard breeding schemes to generate *Ifngr1*^fl/f l^ *Gfap*-Cre^+^ and *Ifngr1*^fl/fl^ *Tie2*-Cre^+^ mice and *Ifngr1*^fl/fl^ littermate controls were used in all experiments. Mice were immunized s.c. with 100 μL of a standard emulsion (Hooke Laboratories) containing complete Freund’s adjuvant and MOG_35-55_ on the upper back and base of the tail. Pertussis toxin (80 ng) (Hooke Laboratories) was injected i.p. on the day of immunization and 2 days later. Mice were monitored daily for clinical signs of disease as follows: 0, no observable signs; 1, limp tail; 2, limp tail and ataxia; 2.5, limp tail and knuckling of at least one limb; 3, paralysis of one limb; 3.5; partial paralysis of one limb and complete paralysis of the other; 4, complete hindlimb paralysis; 4.5, moribund; 5, death.

### qRT-PCR analysis

Total RNA was collected from human brainstem and spinal cord astrocytes (ScienCell) using a RNeasy Kit (QIAGEN) according to manufacturer’s instructions. Reverse transcription and SYBR Green qRT-PCR were performed as previously described (105, 117) for constitutive and immunoproteasome subunits using established primers (118).

### Western blotting

Protein lysates were collected from regional human astrocytes in radioimmunoprecipitation assay (RIPA) buffer (Sigma-Aldrich) supplemented with a protease and phosphatase-3 inhibitor cocktail (Sigma-Aldrich), then 20 μg of protein was resolved on a 4-12% Tris gel and transferred to a PVDF membrane using the Trans-Blot Turbo system (Bio-Rad) according to standard protocols. Membranes were incubated overnight at 4°C in TBS Tween (TBST) plus 5% powdered milk and anti-LMP2 (Abcam; ab184172) or anti-Lys48 (Millipore; 05-1307) and anti-β-actin (ThermoFisher Scientific; MA5-15739) antibodies, washed with TBST 3 times, and then incubated with Alexa Fluor 488, 647, or HRP-conjugated secondary antibodies (ThermoFisher Scientific) for 1 h at room temperature. Membranes were washed with TBST 3 times and imaged using the ChemiDoc MP imaging system (BioRad).

In vitro *ROS and caspase assays*. Human brainstem and spinal cord astrocytes (ScienCell) were seeded in 96-well plates until confluent and treated with media alone or 10 ng/ml IFNγ with or without ONX 0914 for 48 h. ROS were quantified using the DCFDA Cellular ROS Detection Assay Kit (Abcam), caspase activity was quantified using the Caspase-Glo 3/7 Assay Kit (Promega). Cell viability was detected using the CytoPainter Live Cell Labeling Kit (Abcam) and used to normalize ROS and caspase activity. Plates were read on a Victor 3 Multilabel Counter (Perkin Elmer).

### OxyBlot

Human brainstem and spinal cord astrocytes (ScienCell) were seeded in 6-well plates until confluent and treated with media alone or 10 ng/ml IFNγ for 48 h. Protein lysate was isolated in RIPA buffer supplemented with a protease and phosphatase-3 inhibitor cocktail (Sigma-Aldrich). Lysate (20 μg) was then subjected to derivatization according to manufacturer’s instructions (Millipore) and resolved on a 4–12% Tris gel and transferred onto a PVDF transfer membrane (Bio-Rad) using the Trans-Blot Turbo system (Bio-Rad) according to standard protocols. Membranes were incubated overnight at 4°C in TBS Tween (TBST) plus 5% powdered milk and probed with either anti-DNP (Millipore; S7150) or anti-β-actin (ThermoFisher Scientific; MA5-15739) primary antibodies, washed with TBST 3 times, and then incubated with HRP-conjugated secondary antibodies (Millipore) for 1 h at room temperature. Membranes were washed with TBST 3 times, imaged using the ChemiDoc MP imaging system (BioRad), and analyzed as previously described (97).

### Immunoproteasome inhibition

For *in vitro* experiments, the selective immunoproteasome inhibitor ONX 0914 (Cayman Chemical) was dissolved in 0.25% ethanol and used at a working concentration of 50 μM. For *in vivo* immunoproteasome inhibition, ONX 0914 was formulated in an aqueous solution of 14% ethanol in PBS, which was used as a vehicle control, and administered to mice as an i.p. bolus dose of 10 mg/kg after a stabilization or reduction in EAE clinical score for two consecutive days was observed. Seven days after vehicle or ONX 0914 treatment, mice were perfused and CNS tissue was prepared for IHC analysis.

### Immunohistochemistry

Mice were intracardially perfused with PBS followed by 4% PFA, and CNS tissue was removed and fixed in 4% PFA at 4°C for 24 h. Tissue was then cryopreserved in 30% sucrose, and frozen in O.C.T. Compound (Fisher HealthCare). Frozen, sequential transverse sections (12 μm) were slide mounted and stored at −80°C. Tissue sections were blocked with 10% goat serum and 0.1% Triton X-100 (Southern Biotech) for 1 h at room temperature and then incubated with anti-MBP (Abcam; ab7349), -LMP2 (Abcam; ab184172), -PRDX6 (Abcam; ab59543), -Lys48 (Millipore; 05-1307), -GFAP (Invitrogen; 13-0300), -IFNGR1 (ThermoFisher Scientific; 13-1191-82), or -Iba1 (Wako Chemicals; 019-19741) primary antibodies overnight at 4°C. Secondary antibodies conjugated to Alexa Fluor 488 or Alexa Fluor 555 (ThermoFisher Scientific) were applied for 1 h at room temperature. Nuclei were counterstained with DAPI (ThermoFisher Scientific) diluted in PBS. Sections were analyzed using the 20X objective of a confocal microscope LSM 800 (Carl Zeiss). Images shown are representative of at least 4 images taken across two tissue sections at least 100 μm apart per individual mouse. The mean positive area, intensity, and Mander’s coefficient of colocalization were determined by setting thresholds using appropriate isotype control antibodies and quantifying using ImageJ (NIH).

### Statistics

EAE data were analyzed using the nonparametric Mann-Whitney *U* test. Other data were analyzed with parametric tests (2-tailed Student’s *t* test or 2-way ANOVA), with correction for multiple comparisons where appropriate. All statistical analysis was performed using GraphPad Prism Version 7 software (GraphPad Software). A *P* value of less than 0.05 was considered statistically significant.

### Study approval

All experiments were performed in compliance with and under the approval of the Cleveland Clinic Animal Studies Committee.

## Author Contributions

Conceptualization: BCS and JLW; designing research studies: BCS and JLW; conducting experiments: BCS, MS, and JEJ; acquiring data: BCS, MS, JEJ, and MWP; writing the manuscript: BCS, MS, and JLW; editing of the manuscript: BCS, MS, MWP, and JLW; supervision and funding acquisition: JLW.

## Acknowledgements

We thank Dr. Ranjan Dutta for the mRNA expression data, MS tissue and insightful discussion and Dr. Melissa Varrecchia for technical and analytical assistance. We also thank Dr. Bruce Trapp for MS tissue collection, which is supported in part by NINDS R35 NS097303. This work was primarily supported by NIAID K22 AI125466 (JLW).

