## Supplemental Data for "Regional astrocyte interferon-γ signaling regulates immunoproteasome-mediated protection during chronic autoimmunity"

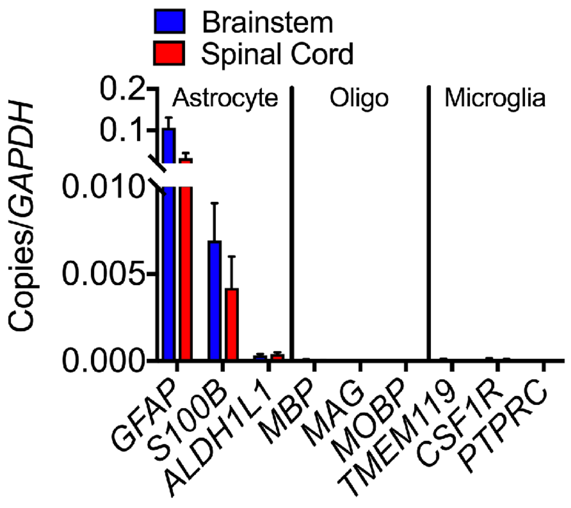


**Figure S1: Confirmation of human astrocyte phenotype.** Human brainstem and spinal cord astrocytes obtained from ScienCell Laboratories were plated in media alone until 80% confluent. RNA was then extracted and qRT-PCR was performed for the indicated transcripts.

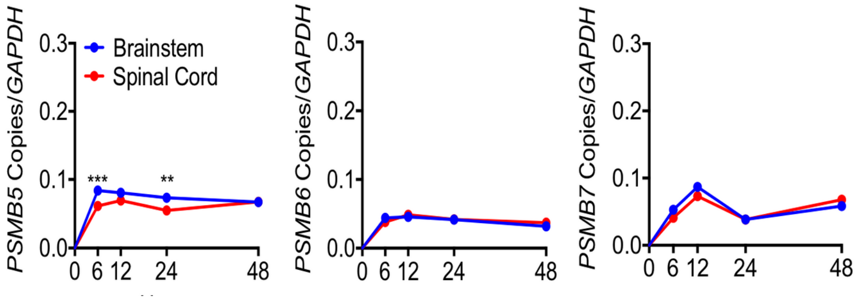


**Figure S2: IFNγ-regulated expression of the constitutive proteasome in regional human astrocytes.** Human brainstem and spinal cord astrocytes were stimulated with 10 ng/ml IFNγ for 0, 6, 12, 24, or 48 h and RNA was collected and analyzed for transcript levels of *PSMB5*, *PSMB6*, and *PSMB7* by qRT-PCR, normalized to copies of *GAPDH*. Data represent the mean ± SEM from 3 independent experiments. **$P < 0.01$, ***$P < 0.001$ between regions by 2-way ANOVA.

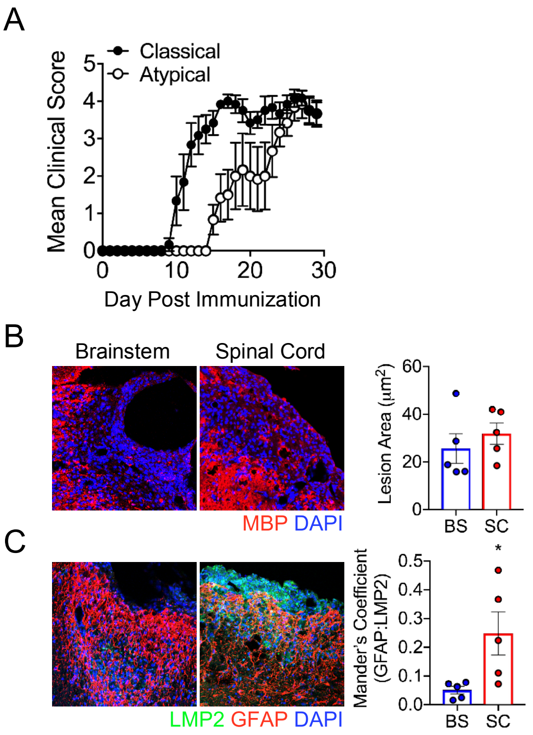


**Figure S3: Regional inflammation and iP expression in *Ifngr1*$^{fl/fl}$*Tie2*-Cre$^{+}$ mice.** EAE was induced in *Ifngr1*$^{fl/fl}$*Tie2*-Cre$^{+}$ mice (*n* = 5) and **(A)** EAE clinical course was monitored. Following 30 days post-immunization, mice were perfused and the CNS was removed and cryopreserved for IHC analysis. White matter tracts of the brain stem and lumbar spinal cord were imaged using confocal microscopy at 20x magnification. (**B**) Tissue sections were labeled for MBP (red) and nuclei were counterstained with DAPI (blue). (**D**) Lesion area was quantified using ImageJ software. **(C)** Tissue sections were labeled for LMP2 (green), GFAP (red), and nuclei were counterstained with DAPI (blue). **(F)** Total LMP2 area and LMP2 colocalized with GFAP were analyzed. *$P$ < 0.05 between regions by 2-tailed Student's *t* test.

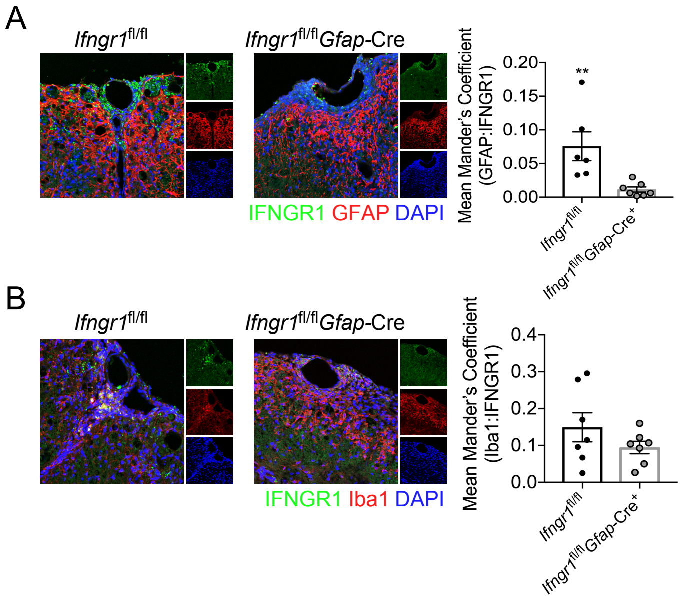


**Figure S4: IFNGR1 deletion in astrocytes of *Ifngr1*$^{fl/fl}$*Gfap*-Cre$^+$ mice. (A)** IHC detection of astrocyte marker GFAP (red) or **(B)** Iba1 (red) and IFNGR1 (green) in the ventral spinal cords of *Ifngr1*$^{fl/fl}$ and *Ifngr1*$^{fl/fl}$*Gfap*-Cre$^+$ mice at day 25 post-EAE induction. Nuclei are shown in blue. Images are representative of at least 4-20x images for each of 7 independent mice per genotype. Colocalization is quantified by Mean Mander's Coefficient using ImageJ software. Data points are representative of individual mice. ***P* < 0.01 between genotypes by 2-tailed Student's *t* test.
